# Recipient-biased competition for a cross-fed nutrient is required for coexistence of microbial mutualists

**DOI:** 10.1101/144220

**Authors:** Alexandra L. McCully, Breah LaSarre, James B. McKinlay

## Abstract

Many mutualistic microbial relationships are based on nutrient cross-feeding. Traditionally, cross-feeding is viewed as being unidirectional from the producer to the recipient. This is likely true when a producer’s metabolic waste, such as fermentation products, provides carbon for a recipient. However, in some cases the cross-fed nutrient holds value for both the producer and the recipient. In such cases, there is potential for nutrient reacquisition by producer cells in a population, leading to competition against recipients. Here we investigate the consequences of inter-partner competition for cross-fed nutrients on mutualism dynamics using an anaerobic coculture pairing fermentative *Escherichia coli* and phototrophic *Rhodopseudomonas palustris*. In this coculture, *E. coli* excretes waste organic acids that provide carbon for *R. palustris*. In return, *R. palustris* cross-feeds *E. coli* ammonium (NH_4_^+^), a valuable nitrogen compound that both species prefer. To explore the potential for inter-partner competition, we first used a kinetic model to simulate cocultures with varied affinities for NH_4_^+^ in each species. The model predicted that inter-partner competition for cross-fed NH_4_^+^ could profoundly impact population dynamics. We then experimentally tested the predictions by culturing mutants lacking NH_4_^+^ transporters in both NH_4_^+^ competition assays and cooperative cocultures. Both theoretical and experimental results indicated that the recipient must have a competitive advantage in acquiring valuable cross-fed NH_4_^+^ to avoid collapse of the mutualism. Thus, the very metabolites that form the basis for cooperative cross-feeding can also be subject to competition between mutualistic partners.

**Significance:** Mutualistic relationships, particularly those based on nutrient cross-feeding, promote stability of diverse ecosystems and drive global biogeochemical cycles. Cross-fed nutrients within these systems can be either waste products valued only by one partner or nutrients that both partners value. Here, we explore how inter-partner competition for a communally-valuable cross-fed nutrient impacts mutualism dynamics. We discovered that mutualism stability necessitates that the recipient have a competitive advantage against the producer in obtaining the cross-fed nutrient. We propose that the requirement for recipient-biased competition is a general rule for mutualistic coexistence based on the transfer of communally valuable resources, microbial or otherwise.

## Introduction

Mutualistic cross-feeding of resources between microbes can have broad impacts ranging from influencing host health (1, 2) to driving global biogeochemical cycles (3–6). Cross-fed metabolites are often regarded as nutrients due to the value they provide to a dependent partner, the recipient. However, for the partner producing the nutrient, the producer, a cross-fed nutrient’s value can vary. On one extreme, the cross-fed metabolite is valued by the recipient but not the producer, as is the case for fermentative waste products (7–10). In other cases, a cross-fed metabolite holds value for both the recipient and the producer, as is the case for vitamin B_12_ (6, 11, 12) and ammonium (NH_4_^+^) (13, 14). Such communally-valuable cross-fed nutrients are subject to partial privatization (15), wherein the producer has mechanisms to retain a portion of the nutrient pool for itself. While most mutualism cross-feeding studies only consider unidirectional metabolite transfer from producer to recipient, we wondered whether these mechanisms for partial privatization could lead to competition between partner populations for communally-valuable cross-fed nutrients. It seems likely that such competition could influence mutualism stability, as is known to be the case for competition for exogenous limiting resources (8, 16–19). To the best of our knowledge inter-partner competition for cross-fed nutrients and its impact on mutualism dynamics have never been investigated.

One example of cross-feeding that could involve competition between mutualistic partners is NH_4_^+^ excretion by N_2_-fixing bacteria (Fig. 1A), called N_2_-fixers (13, 14). During N_2_ fixation, the enzyme nitrogenase converts N_2_ gas into two NH_3_ (20). In an aqueous environment, NH_3_ is in equilibrium with NH_4_^+^. At neutral pH, NH_4_^+^ is the predominant form but small amounts of NH_3_ can potentially leave the cell by diffusion across the membrane (21) (Fig. 1B). This inherent ‘leakiness’ for NH_3_ likely fosters NH_4_^+^ cross-feeding, as extracellular NH_3_ is available to neighboring microbes. Importantly, these neighbors can include clonal N_2_-fixers, as NH_3_/NH_4_^+^ is a preferred nitrogen source for most microbes. At concentrations above 20 μM, NH_3_ can be acquired by passive diffusion; below 20 μM, NH_4_^+^ is bound and transported as NH_3_ by AmtB transporters (Fig. 1B) (22). AmtB-like transporters are conserved throughout all domains of life (23). There is growing evidence that AmtB is used by N_2_-fixers to recapture NH_3_ lost by passive diffusion, as ΔAmtB mutants accumulate NH_4_^+^ in culture supernatants whereas wild-type strains do not (24–26). Thus, during NH_4_^+^ cross-feeding, AmtB likely facilitates both NH_4_^+^ acquisition by the mutualistic partner and recapture of NH_4_^+^ by the N_2_-fixer.

**Fig. 1.**
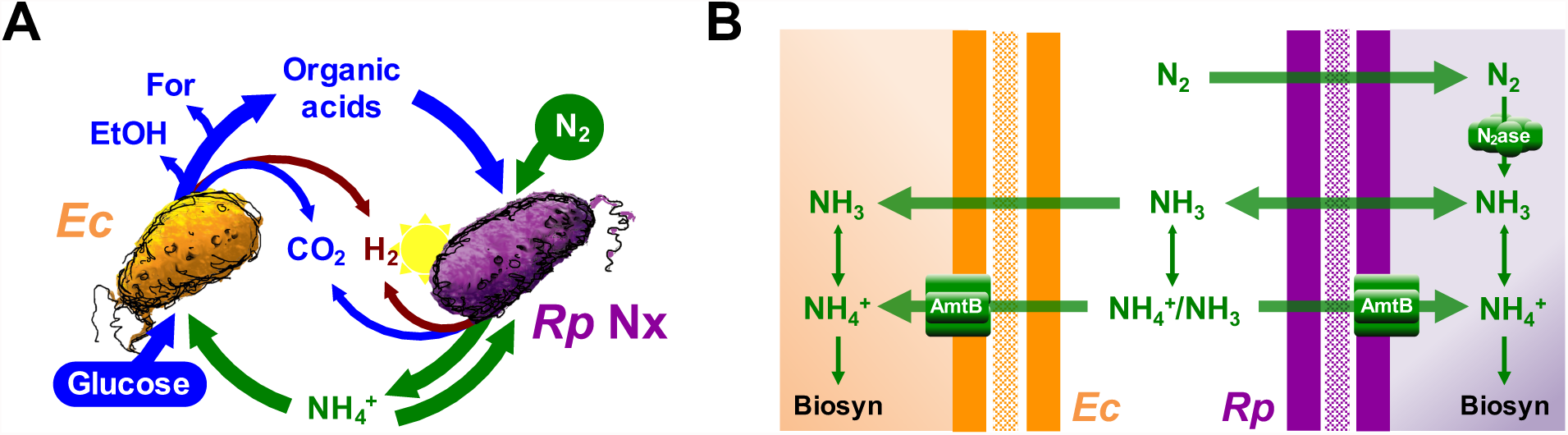
Mechanisms of NH_4_^+^ transfer within an obligate bacterial mutualism based on cross-feeding of essential nutrients. (**A**) *Escherichia coli* (*Ec*) anaerobically ferments glucose into organic acids, supplying *Rhodopseudomonas palustris* Nx (*Rp* Nx) with essential carbon. *R. palustris* Nx fixes N_2_ gas and excretes NH_4_^+^, supplying *E. coli* with essential nitrogen. For, formate; EtOH, ethanol. (**B**) NH_4_^+^ can be passively lost from cells as NH_3_. Both species encode high-affinity NH_4_^+^ transporters, AmtB, that facilitate NH_4_^+^ uptake. NH_4_^+^ is the predominant form at neutral pH, as indicated by an enlarged arrow head on double-sided arrows.

Assessing the effects of inter-partner competition for a cross-fed nutrient would require a level of experimental control not possible in most natural settings. However, synthetic microbial communities, or cocultures, are well-suited to address such questions (27–29). We previously developed a bacterial coculture that features cross-feeding of waste products (organic acids) from *Escherichia coli*, and a communally-valuable nutrient (NH_4_^+^) from *Rhodopseudomonas palustris* Nx (Fig. 1A) (26). Here, using both a kinetic model and genetic manipulation to alter the affinity of each species in the coculture for NH_4_^+^, we demonstrate that inter-partner competition for cross-fed NH_4_^+^ plays a direct role in maintaining coexistence. Specifically, insufficient competition by *E. coli* for NH_4_^+^ resulted in a collapse of the mutualism. Mutualism collapse could be delayed or potentially avoided through higher net NH_4_^+^ excretion by *R. palustris* or increased *E. coli* population size. Our results suggest that, as a general rule, competition for a cross-fed nutrient in an obligate mutualism must be biased in favor of the recipient to avoid mutualism collapse and the potential extinction of both species.

## Results

### Competition for cross-fed NH_4_^+^ is predicted to shape mutualism population dynamics

Within our coculture (Fig. 1A), *E. coli* (*Ec*) ferments sugars into waste organic acids, providing essential carbon and electrons to *R. palustris* (*Rp*) Nx. *R. palustris* Nx is genetically engineered to excrete low micromolar amounts of NH_4_^+^, providing essential nitrogen for *E. coli* (26). NH_4_^+^ excretion by *R. palustris* Nx is due to mutation of NifA, the master transcriptional regulator of nitrogenase, which results in constitutive nitrogenase activity even in the presence of normally inhibitory NH_4_^+^ (30). In contrast to organic acids, which are only useful to *R. palustris*, NH_4_^+^ produced by *R. palustris* Nx is essential for the growth of both species; *R. palustris* uses some NH_4_^+^ for its own biosynthesis and excretes the rest, which serves as the nitrogen source for *E. coli.* However, *R. palustris* Nx can also take up NH_4_^+^ (30). Thus, we hypothesized that competition for cross-fed NH_4_^+^ between the *R. palustris* Nx producer population and the *E. coli* recipient population could influence mutualism dynamics.

We first explored whether competition for cross-fed NH_4_^+^ could affect the mutualism using SyFFoN, a mathematical model describing our coculture (26, 31). SyFFoN simulates population and metabolic dynamics in batch cocultures using Monod equations with experimentally-determined parameter values. As previous versions described NH_4_^+^ uptake kinetics only for *E. coli* (26, 31), we amended SyFFoN to include both an *R. palustris* NH_4_^+^ uptake affinity (K_m_) and higher *R. palustris* maximum growth rate (μ_MAX_) when NH_4_^+^ is used (SI Appendix Table S1). We then simulated batch cocultures wherein the relative affinity for NH_4_^+^ varied between the two species (Fig. 2). The model predicted that coexistence is maintained when the *R. palustris* affinity for NH_4_^+^ is low relative to that of *E. coli* (*Rp*:*Ec* < 1); sufficient N_2_ is converted to NH_4_^+^ to support *R. palustris* growth and enough NH_4_^+^ is cross-fed to support *E. coli* growth. In contrast, when the *R. palustris* affinity for NH_4_^+^ is high relative to that of *E. coli* (*Rp*:*Ec* > 1), *E. coli* growth is no longer supported because *E. coli* cannot compete for excreted NH_4_^+^. However, high *R. palustris* cell densities were still predicted (Fig. 2) due to persistent, low-level organic acid cross-feeding stemming from *E. coli* maintenance metabolism, which can support *R. palustris* growth even when *E. coli* is not growing (31).

**Fig. 2.**
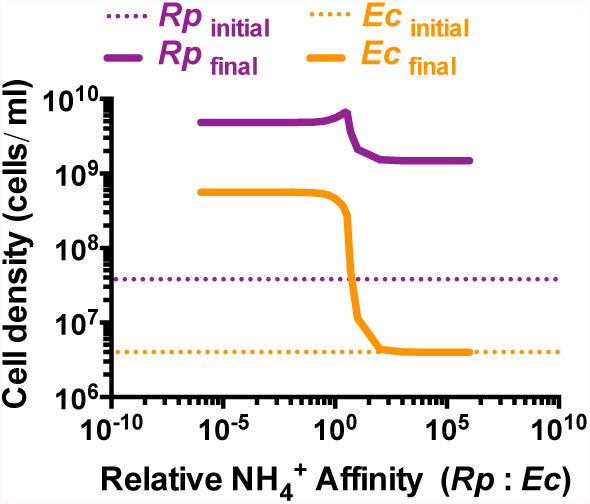
Simulations suggest that *E. coli* must have a competitive advantage for NH_4_^+^ acquisition relative to *R. palustris* to support mutualistic growth. Final cell densities (solid lines) of *R. palustris* (*Rp*, purple) and *E. coli* (*Ec*, orange) after 300 h in simulated batch cultures for a range of relative NH_4_^+^ affinities. Initial cell densities are indicated by dotted lines. Relative NH_4_^+^ affinity values represent the relative *E. coli* K_m_ for NH_4_^+^ (K_A_) divided by that of *R. palustris* (K_AR_).

### Genetic disruption of AmtB NH_4_^+^ transporters affects relative affinities for NH_4_^+^

Bacterial cells generally acquire NH_4_^+^ through two mechanisms: passive diffusion of NH_3_, or active uptake by AmtB transporters (Fig. 1B). We hypothesized that deleting *amtB* genes in either species would result in a lower affinity for NH_4_^+^ in that species and thus could be used to test how relative NH_4_^+^ affinity impacts coculture dynamics. We generated ΔAmtB mutants of both *E. coli* and *R. palustris* and first characterized the effect of the mutations in monoculture. Deletion of *amtB* in *E. coli* had no effect on growth or fermentation profiles when NH_4_Cl was in excess (SI Appendix Fig. S1), consistent with previous observations where ΔAmtB growth defects were only apparent at NH_4_^+^ concentrations below 20 μM (22). In *R. palustris* ΔAmtB monocultures with N_2_ as the nitrogen source, growth trends were equivalent to those of the parent strain; however, *R. palustris* ΔAmtB excreted more NH_4_^+^ than the parent strain and about a third of that excreted by *R. palustris* Nx (SI Appendix Fig. S1C and D). In line with our hypothesis, NH_4_^+^ excretion by *R. palustris* ΔAmtB could be due to a decreased ability to reacquire NH_4_^+^ lost by diffusion, resulting in increased net NH_4_^+^ excretion. Alternatively, we considered that NH_4_^+^ excretion by *R. palustris* ΔAmtB could be due to improper nitrogenase regulation. In several other N_2_-fixers, proper nitrogenase regulation requires AmtB, for example to induce post-translational nitrogenase inhibition (switch-off) in response to NH_4_^+^ (25, 32). We tested whether *R. palustris* ΔAmtB exhibits NH_4_^+^-induced switch-off by adding NH_4_Cl to exponentially growing cultures and measuring H_2_ production, an obligate product of the nitrogenase reaction (33), as a proxy for nitrogenase activity. Upon adding NH_4_Cl, H_2_ production stopped in *R. palustris* ΔAmtB cultures. In contrast, H_2_ production only slowed slightly in *R. palustris* Nx cultures (SI Appendix Fig. S2), consistent with previous observations for NifA* strains (34, 35). Additionally, like the parent strain, *R. palustris* ΔAmtB did not produce H_2_ when grown with NH_4_^+^, unlike *R. palustris* Nx (SI Appendix Fig. S3). These observations demonstrate that *R. palustris* ΔAmtB is competent for NH_4_^+^-induced nitrogenase repression, and thus NH_4_^+^ excretion by *R. palustris* ΔAmtB is likely due to a poor ability to reacquire NH_4_^+^ lost by diffusion.

To test our hypothesis that deleting *amtB* would lower cellular affinity for NH_4_^+^, we directly competed all possible *E. coli* and *R. palustris* strain combinations in competition assays where ample carbon was available for each species but the NH_4_^+^ concentration was kept low; specifically, a small amount of NH_4_^+^ was added every hour to bring the final NH_4_^+^ concentration to 5 μM (Fig. 3). In this competition assay, the species that is more competitive for NH_4_^+^ should reach a higher cell density than the other species. In all cases, WT *E. coli* was more competitive for NH_4_^+^ than *R. palustris*. However, each *R. palustris* strain was able to outcompete *E. coli* ΔAmtB (Fig. 3), even though the *R. palustris* maximum growth rate is 4.6-times slower than that of *E. coli* (SI Appendix Fig. S1). Even *R. palustris* strains lacking AmtB outcompeted *E. coli* ΔAmtB (Fig. 3), indicating that *R. palustris* has a higher affinity for NH_4_^+^ than *E. coli* independent of AmtB. These data confirmed that deletion of *amtB* was an effective means by which to lower the relative affinity for NH_4_^+^ in each mutualistic partner.

**Fig. 3.**
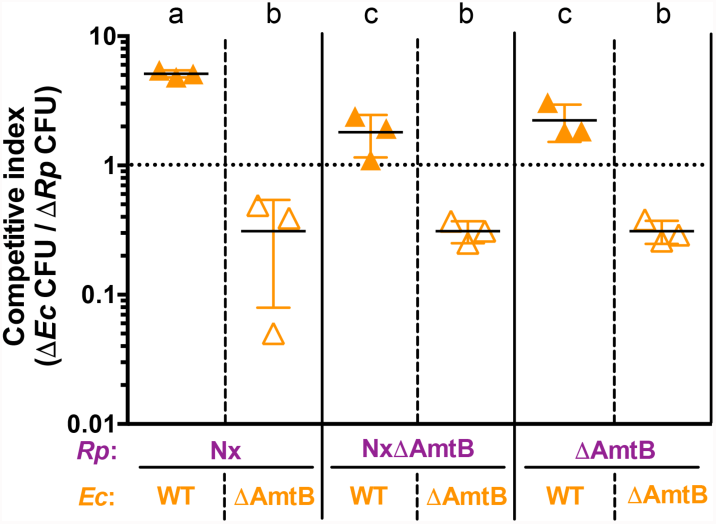
AmtB is important for competitive NH_4_^+^ acquisition. Competitive indices for *E. coli* after 96 h in NH_4_^+^-limited competition assay cocultures. Cocultures were inoculated with *E. coli* and *R. palustris* at equivalent cell densities with excess carbon available for both species (25 mM glucose for *E. coli* and 20 mM sodium acetate for *R. palustris*). NH_4_^+^ was added to cocultures to a final concentration of 0.5 μM every hour for 96 h. The dotted line indicates a competitive index value of 1, where both species are equally competitive for NH_4_^+^. Filled triangles, WT *E. coli*; open triangles *E. coli* ΔAmtB. Error bars indicate SD, n=3. Different letters indicate statistical differences between *E. coli* competitive index values, p < 0.05, determined by one-way ANOVA with Tukey’s multiple comparisons post test.

### Altering relative NH_4_^+^ affinities affects mutualistic partner frequencies

We then examined how relative affinities for NH_4_^+^ influenced mutualism dynamics by comparing the growth trends of cocultures containing either WT *E. coli* or *E. coli* ΔAmtB, paired with either *R. palustris* ΔAmtB, *R. palustris* Nx, or *R. palustris* NxΔAmtB, the latter of which we previously determined to exhibit 3-fold higher NH_4_^+^- excretion levels than the Nx strain in monoculture (26). For each *R. palustris* partner, cocultures with *E. coli* ΔAmtB grew slower than cocultures with WT *E. coli* (Fig. 4A,B). *E. coli* ΔAmtB also constituted a lower percentage of the population and achieved lower cell densities compared to WT *E. coli* when paired with the same *R. palustris* strain (Fig. 4C). These lower frequencies were consistent with the competitive disadvantage of *E. coli* ΔAmtB for excreted NH_4_^+^ (Fig. 3).

**Fig. 4.**
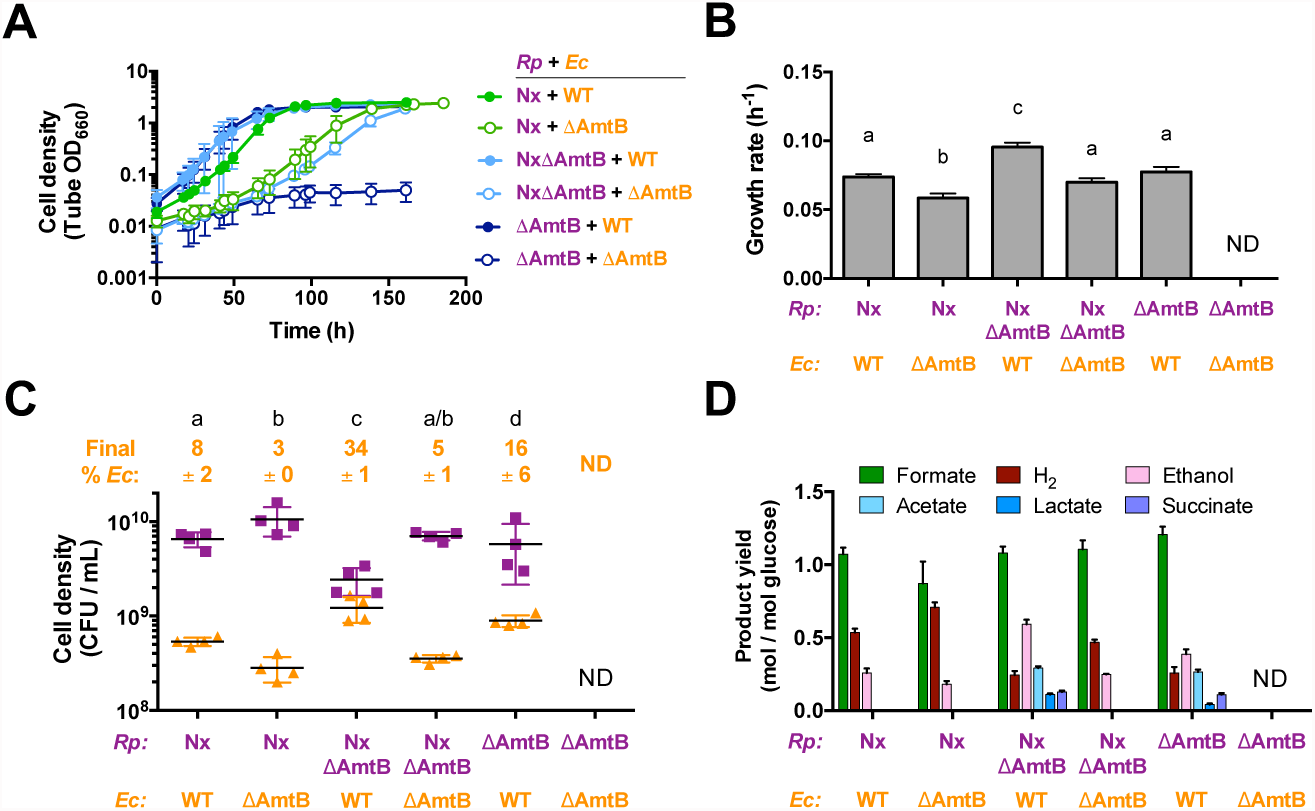
NH_4_^+^ transporters influence population and metabolic trends of both partners in coculture. Growth curves (**A**), growth rates (**B**), final cell densities after one culturing (**C**), and fermentation product yields (**D**) from cocultures of all combinations of mutants lacking AmtB in each species. Final cell densities and fermentation product yields were taken after one week, within 24 h into stationary phase. ND, not determined. Error bars indicate SD, n=4. Different letters indicate statistical differences, p < 0.05, determined by one-way ANOVA with Tukey’s multiple comparisons post test.

For *R. palustris* strains lacking AmtB, the effects on population trends varied. Consistent with our previous work, *R. palustris* NxΔAmtB supported higher WT *E. coli* percentages and cell densities (Fig. 4C) (26). With high NH_4_^+^ excretion levels from *R. palustris* NxΔAmtB, faster *E. coli* growth leads to rapid organic acid accumulation, which acidifies the environment, inhibits *R. palustris* growth, and leaves organic acids unconsumed (Fig. 4D) (26). Surprisingly, although *R. palustris* ΔAmtB excreted less NH_4_^+^ than *R. palustris* Nx in monoculture, *R. palustris* ΔAmtB supported a higher WT *E. coli* population in coculture and consumable organic acids accumulated (Fig. 4C, D). These trends resemble those from cocultures with *R. palustris* NxΔAmtB (Fig. 4C, D), which has a high level of NH_4_^+^ excretion (SI Appendix Fig. S1D). Unlike Nx strains, which have constitutive nitrogenase activity due to a mutation in the transcriptional activator NifA (30), *R. palustris* ΔAmtB has WT NifA. Thus, *R. palustris* ΔAmtB can likely still regulate nitrogenase expression, and thereby its activity, in response to nitrogen starvation. We hypothesized that in coculture with WT *E. coli*, *R. palustris* ΔAmtB might experience heightened nitrogen starvation, as NH_4_^+^ consumption by WT *E. coli* would limit NH_4_^+^ reacquisition by *R. palustris* ΔAmtB (in an *R. palustris* ΔAmtB monoculture any lost NH_4_^+^ would remain available to *R. palustris*). We therefore tested whether coculture conditions stimulated higher nitrogenase activity using an acetylene reduction assay. In agreement with our hypothesis, *R. palustris* ΔAmtB had increased nitrogenase activity in coculture conditions compared to monocultures, whereas *R. palustris* Nx, which exhibits constitutive nitrogenase activity, showed similar levels in both conditions (SI Appendix Fig. S4). Thus, the relatively high WT *E. coli* population in coculture with *R. palustris* ΔAmtB is likely due to both the competitive advantage for acquiring NH_4_^+^ over *R. palustris* ΔAmtB (Fig. 3) and higher NH_4_^+^ cross-feeding levels due to increased nitrogenase activity.

### *E. coli* must have a competitive advantage for NH_4_^+^ acquisition to avoid mutualism collapse

We were surprised to observe that cocultures of *R. palustris* ΔAmtB paired with *E. coli* ΔAmtB showed little growth when started from a single colony of each species (Fig. 4A), a method that we routinely use to initiate cocultures (26, 31). We reasoned that the higher *R. palustris* ΔAmtB affinity for NH_4_^+^ relative to *E. coli* ΔAmtB (Fig. 3) likely led to community collapse as predicted by SyFFoN (Fig. 2). Even though SyFFoN had predicted *R. palustris* growth when outcompeting *E. coli* for NH_4_^+^ (Fig. 2), SyFFoN likely underestimates the time required to achieve these densities, if they would be achieved at all, as SyFFoN does not take into account cell death, which is known to occur when *E. coli* growth is prevented (31). Consistent with the hypothesis that poor coculture growth was due to a competitive disadvantage of *E. coli* ΔAmtB for NH_4_^+^, SyFFoN simulations indicated that starting with a larger *E. coli* ΔAmtB population would increase the probability that any given *E. coli* cell would acquire NH_4_^+^ versus *R. palustris* and thereby overcome the competitive disadvantage of *E. coli* ΔAmtB for NH_4_^+^ (SI Appendix Fig. S5). Indeed, we observed greater growth of both species when cocultures were inoculated with equal or higher relative densities of *E. coli* ΔAmtB versus *R. palustris* ΔAmtB (SI Appendix Fig. S5).

The explanation that mutualism collapse was due to a competitive advantage of *R. palustris* ΔAmtB over *E. coli* ΔAmtB for NH_4_^+^ called into question why cocultures pairing *E. coli* ΔAmtB with either *R. palustris* Nx or *R. palustris* NxΔAmtB did not collapse as well (Fig. 4), given that in all of these pairings *E. coli* ΔAmtB is at competitive disadvantage (Fig. 3). We hypothesized that a relatively high NH_4_^+^ excretion level by these latter *R. palustris* strains (SI Appendix Fig. S1D) could compensate for a low *E. coli* NH_4_^+^ affinity. To explore this hypothesis we simulated cocultures with the *R. palustris* affinity for NH_4_^+^ set high relative to that of *E. coli* (*Rp*:*Ec* = 1000) and varied the *R. palustris* NH_4_^+^ excretion level (Fig. 5). Indeed, increasing *R. palustris* NH_4_^+^ excretion was predicted to overcome a low *E. coli* affinity for NH_4_^+^ and support growth of both species (Fig. 5). The only exception was at the highest levels of NH_4_^+^ excretion, where *R. palustris* growth was predicted to be inhibited due to rapid *E. coli* growth and subsequent accumulation of organic acids that acidify the environment (Fig. 5) (26). These simulations suggested that *R. palustris* Nx and NxΔAmtB supported coculture growth with *E. coli* ΔAmtB due to higher NH_4_^+^ excretion levels (SI Appendix Fig. S1D), whereas a combination of low NH_4_^+^ excretion by *R. palustris* ΔAmtB (SI Appendix Fig. S1D) and a low affinity for NH_4_^+^ by *E. coli* ΔAmtB led to collapse of the mutualism in this pairing.

**Fig. 5.**
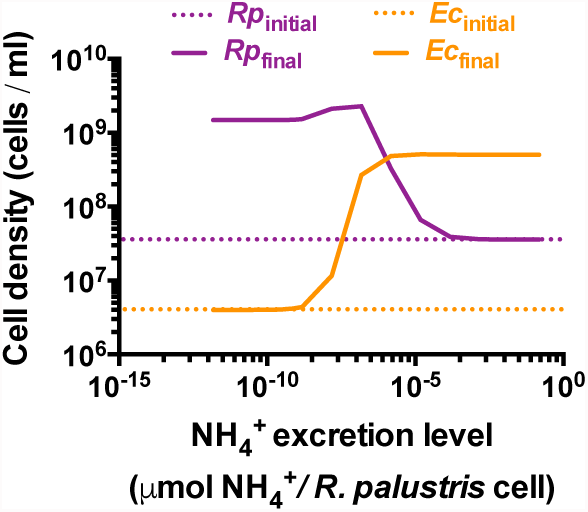
Higher *R. palustris* NH_4_^+^ excretion levels are predicted to compensate for a low *E. coli* NH_4_^+^ affinity. 300 h batch cultures were simulated with a relative *R. palustris* : *E. coli* (*Ec* : *Rp*) K_m_ value for NH_4_^+^ of 0.001 over different *R. palustris* NH_4_^+^ excretion levels (R_A_). Final cell densities, solid lines; initial cell densities, dotted lines.

So far, we had only considered the effect of severe discrepancies in NH_4_^+^ affinities between the two species (e.g., 1000-fold difference in K_m_ values in our simulations) as a mechanism leading to coculture collapse within the time period of a single culturing. However, we wondered if a subtle discrepancy in NH_4_^+^ affinities could lead to coculture collapse if given more time. We therefore simulated serial transfers of cocultures with partners having different relative NH_4_^+^ affinities (Fig. 6A, B). At equivalent NH_4_^+^ affinities (Fig. 6A), both species were predicted to be maintained over serial transfers. However, when the relative affinities approached a threshold (relative *Rp*:*Ec* = 2.75), cell densities of both species were predicted to decrease over serial transfers (Fig. 6B). This decline in coculture growth is due to *E. coli* being slowly but progressively outcompeted for NH_4_^+^ by *R. palustris*. As the difference between the *R. palustris* and *E. coli* populations expands, *R. palustris* cells have a greater chance of acquiring NH_4_^+^ than the smaller *E. coli* population, further starving *E. coli* and simultaneously cutting off *R. palustris* from its supply of organic acids from *E. coli.*

**Fig. 6.**
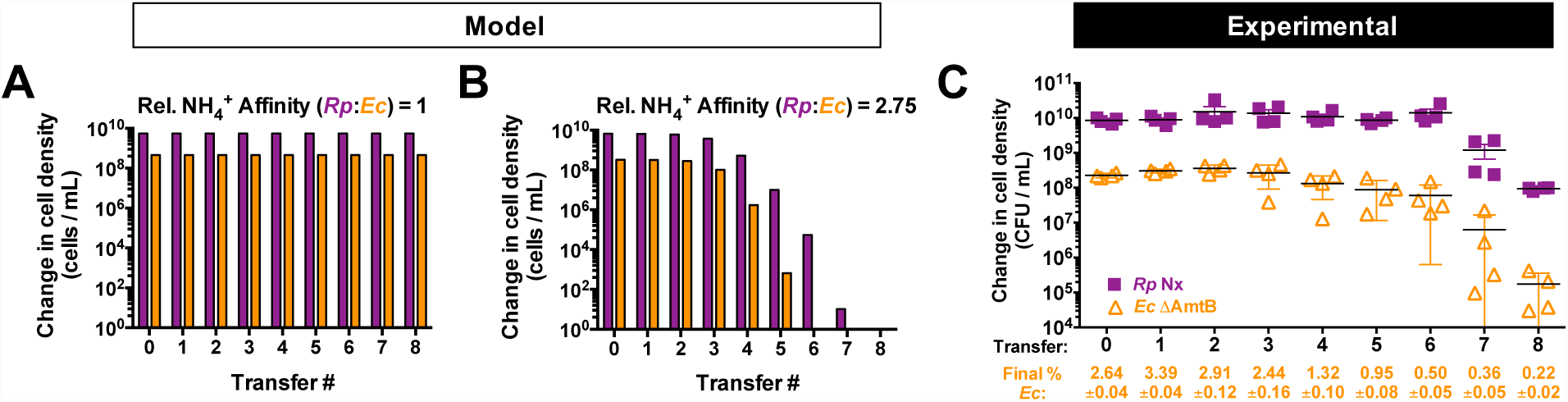
A low *E. coli* NH_4_^+^ affinity results in coculture collapse over serial transfers when paired with *R. palustris* Nx. (**A**,**B**) 300 h batch cultures were simulated and serial transferred used a 1% inoculum based on the cell density at 300 h for the previous culture. Relative NH_4_^+^ affinity values represent the relative *E. coli* K_m_ for NH_4_^+^ (K_A_) divided by that of *R. palustris* (K_AR_). (**C**) Final cell densities of *R. palustris* Nx and *E. coli* ΔAmtB of cocultures grown for one week, less than 24 h into stationary phase. A 1% inoculum was used for each subsequent serial transfer. Error bars indicate SD, n=4. Final *E. coli* cell percentages +/− SD for each transfer are shown.

The above prediction prompted us to investigate if cocultures pairing *R. palustris* Nx with *E. coli* ΔAmtB were stable through serial transfers. We focused on cocultures with *R. palustris* Nx rather than *R. palustris* NxΔAmtB because *R. palustris* Nx has AmtB and would therefore be most likely to outcompete *E. coli* ΔAmtB. Strikingly, after eight serial transfers of cocultures pairing *R. palustris* Nx with *E. coli* ΔAmtB we observed coculture collapse (Fig. 6C). This observation is in stark contrast to cocultures of *R. palustris* Nx paired with WT *E. coli*, which we have serially transferred for over 100 times with no extinction events (unpublished data). These results indicate that the recipient population must have a competitive advantage for a cross-fed nutrient versus the producer population to avoid mutualism collapse.

## Discussion

Here we demonstrate that mutualistic partners can compete for a cross-fed nutrient upon which the mutualistic interaction is based, in this case NH_4_^+^. This competition can impact partner frequencies and mutualism stability. Efficient nutrient reacquisition by the producer can render nutrient excretion levels insufficient for cooperative growth, starving the recipient and leading to tragedy of the commons (36). Conversely, recipient-biased competition for a cross-fed nutrient drives cooperative directionality in nutrient exchange and thereby promotes mutualism stability. One implication of these results is that inter-partner competition can influence the level of resource privatization. Within microbial interdependencies, partial privatization has primarily been thought to depend on mechanisms used by the producer to retain a portion of a communally-valuable resource (15). Our data indicate that for excreted resources having a transient availability to both mutualists, recipient acquisition mechanisms can also influence the level of producer privatization, as the competition impacts how much of a cross-fed resource will be shared versus re-acquired. In effect, recipient-biased competition avoids tragedy of the commons by enforcing partial privatization of a communally-valuable resource. The importance of the recipient having the upper hand in inter-partner competition likely applies to other synthetic cocultures and natural microbial mutualisms that are based on the cross-feeding of communally-valuable nutrients, including amino acids (37, 38) and vitamin B12 (6, 11). The same rule could also apply to inter-kingdom and non-microbial examples of cross-feeding (e.g., plants and pollinators, nutrient transfer between plants and bacteria or fungi (39)) and cooperative feeding (e.g., honeyguide bird and human harvesting of bee hives (40), cooperative hunting between grouper fish and moray eels (41)). In such cases, increased privatization of a cross-fed or shared resource, for example through producer-biased competition, could threaten the mutualism upon which both species depend (15, 39, 42).

In our system, AmtB transporters were crucial determinants of inter-partner competition for NH_4_^+^. We were intrigued to find that when both species lacked AmtB, *R. palustris* out-competed *E. coli* for NH_4_^+^ (Fig. 5), enough so to collapse the mutualism within a single culturing (Fig. 3). Whether by maximizing NH_4_^+^ retention or re-acquisition, *R. palustris*, and perhaps other N_2_-fixers, might have additional mechanisms aside from AmtB to minimize loss of NH_4_^+^ as NH_3_. These mechanisms could include a relatively low internal pH to favor NH_4_^+^ over NH_3_, negatively-charged surface features, or relatively high affinities by NH_4_^+^-assimilating enzymes such as glutamine synthetase. There are several reasons why it would be beneficial for N_2_-fixers to minimize NH_4_^+^ loss. First, N_2_ fixation is expensive, both in terms of the enzymes involved (43) and the reaction itself, costing 16 ATP to convert one N_2_ into two NH_3_ (33). Passive loss of NH_3_ would only add to this cost, as more N_2_ would have be fixed to compensate. Second, loss of NH_4_^+^ could benefit nearby microbes competing against an N_2_-fixer for separate limiting nutrients (14, 44). The possibility that N_2_-fixers could have a superior ability to retain or acquire NH_4_^+^ independently of AmtB is not farfetched. Bacteria are known to exhibit differential abilities to compete for nutrients. For example, iron acquisition commonly involves iron-binding siderophores, but siderophores can be chemically distinct and thereby differ in their affinity for iron (45). Strategies to utilize siderophores as a shared resource are also numerous, leading to different cooperative or competitive outcomes in microbial communities (45, 46). One must consider that additional mechanisms for acquiring NH_4_^+^ beyond AmtB might likewise exist. As our results have raised the potential for inter-partner competition for cross-fed resources themselves, understanding the physiological mechanisms that confer competitive advantages for nutrient acquisition between species will undoubtedly aid in describing the interplay between competition and cooperation within mutualisms.

## Materials and Methods

### Strains and growth conditions

Strains, plasmids, and primers are listed in SI Appendix Table S2. All *R. palustris* strains contained *ΔuppE* and *ΔhupS* mutations to facilitate accurate colony forming unit (CFU) measurements by preventing cell aggregation (47) and to prevent H_2_ uptake, respectively. *E. coli* was cultivated on Luria-Burtani (LB) agar and *R. palustris* on defined mineral (PM) (48) agar with 10 mM succinate. (NH_4_)_2_SO_4_ was omitted from PM agar for determining *R. palustris* CFUs. Monocultures and cocultures were grown in 10-mL of defined M9-derived coculture medium (MDC) (26) in 27-mL anaerobic test tubes. To make the medium anaerobic, MDC was bubbled with N_2_, then tubes were sealed with rubber stoppers and aluminum crimps, and then autoclaved. After autoclaving, MDC was supplemented with cation solution (1 % v/v; 100 mM MgSO_4_ and 10 mM CaCl_2_) and glucose (25 mM), unless indicated otherwise. *E. coli* monocultures were also supplemented with 15mM NH_4_Cl. All cultures were grown at 30°C laying horizontally under a 60 W incandescent bulb with shaking at 150 rpm. Starter cocultures were inoculated with 200 μL MDC containing a suspension of a single colony of each species. Test cocultures were inoculated using a 1% inoculum from starter cocultures. Serial transfers were also inoculated with a 1% inoculum. Kanamycin and gentamycin were added to a final concentration of 100 μg/ml for *R. palustris* and 15 μg/ml for *E. coli* where appropriate.

### Generation of *R. palustris* mutants

*R. palustris* mutants were derived from wild-type CGA009 (49). Generation of strains CGA4004, CGA4005, and CGA4021 was described previously (26). For generation of strain CGA4026 (*R. palustris* ΔAmtB) the WT *nifA* gene was amplified using primers JBM1 and JBM2, digested with XbaI and BamHI, and ligated into plasmid pJQ200SK to make pJQnifA16. This suicide vector was then introduced into CGA4021 by conjugation, and sequential selection and screening was performed as described (50) to replace *nifA\** with WT *nifA*. Reintroduction of the WT *nifA* gene was confirmed by PCR and sequencing.

### Generation of the *E. coli* ΔAmtB mutant

P1 transduction (51) was used to introduce Δ*amtB::Km* from the Keio collection strain JW0441-1 (52) into MG1655. The Δ*amtB::Km* genotype of kanamycin-resistant colonies was confirmed by PCR and sequencing.

### Analytical procedures

Cell density was assayed by optical density at 660 nm (OD_660_) using a Genesys 20 visible spectrophotometer (Thermo-Fisher, Waltham, MA, USA). Growth curve readings were taken in culture tubes without sampling (i.e., Tube OD_660_). Specific growth rates were determined using readings between 0.1-1.0 OD_660_ where there is linear correlation between cell density and OD_660_. Final OD_660_ measurements were taken in cuvettes and samples were diluted into the linear range as necessary. H_2_ was quantified using a Shimadzu (Kyoto, Japan) gas chromatograph (GC) with a thermal conductivity detector as described (53). Glucose, organic acids, formate and ethanol were quantified using a Shimadzu high-performance liquid chromatograph (HPLC) as described (54). NH_4_^+^ was quantified using an indophenol colorimetric assay as described (26).

### Nitrogenase activity

Nitrogenase activity was measured using an acetylene reduction assay (43). Cells from 10-mL cultures were harvested and resuspended in 10-mL fresh MDC medium in 27-mL sealed tubes pre-flushed with argon gas. Suspensions were incubated in light for 1 h at 30°C to recover. Then, 250 μl of 100% acetylene gas was injected into the headspace to initiate the assay, and ethylene production was measured over time by gas chromatography as described (43). Ethylene levels were normalized to total *R. palustris* CFUs in the 10-ml volume.

### NH_4_^+^ competition assay

Fed-batch cultures were performed in custom anaerobic 75-ml serum vials with side sampling ports. Each vial contained a stir bar and 30-mL of MDC, and was sealed at both ends with rubber stoppers and aluminum crimps. Each vial was supplemented with 25 mM glucose, 1 % v/v cation solution and 20 mM sodium acetate. Starter monocultures of each species were grown to equivalent CFUs/mL in MDC tubes containing limiting nutrients (3 mM sodium acetate for *R. palustris* and 1.5 mM NH_4_Cl for *E. coli*), and 1 mL of each species was inoculated into the serum vials. These competition cocultures were incubated at 30°C under a 60 W incandescent bulb with stirring at 200 rpm (Thermo Scientific) for 96 h. Each serum vial was constantly flushed with Ar to maintain anaerobic conditions. NH_4_Cl was fed from a 500 μM NH_4_Cl stock using a peristaltic pump (Watson-Marlow) on an automatic timer (Intermatic DT620) at a rate of 0.33 mL/min once an hour for a final concentration of ~ 5 μM upon each addition. Samples were taken at 0 and 96 h for quantification of CFUs.

### Mathematical modeling

A Monod model describing bi-directional cross-feeding in batch cultures, called SyFFoN_v3 (Syntrophy between Fermenter and Fixer of Nitrogen), was modified from our previous model (31) to allow for competition between *E. coli* and *R. palustris* for NH_4_^+^ as follows: (i) an equation for *R. palustris* growth rate on NH_4_^+^ was added to boost the *R. palustris* growth rate when acquiring NH_4_^+^ and (ii) the ability for *R. palustris* to consume NH_4_^+^ was added along with a K_m_ of *R. palustris* for NH_4_^+^ (K_AR_). Equations and default parameter values are in the SI Appendix and Table S1. SyFFoN_v3 runs in R studio and is available for download at: https://github.com/McKinlab/Coculture-Mutualism.

## Acknowledgments

We thank Richard Phillips (Indiana University) for providing equipment for the NH_4_^+^ competition assay. We also thank Jay Lennon (Indiana University) for helpful discussions on the manuscript. This work was supported in part by the U.S. Department of Energy, Office of Science, Office of Biological and Environmental Research, under Award Number DE-SC0008131, by the U.S. Army Research Office, grant W911NF-14-1-0411, and by the Indiana University College of Arts and Sciences.

## SI Appendix

### SyFFoN_v3 description

Equations 1 – 4 were used to describe *E. coli* and *R. palustris* growth rates:

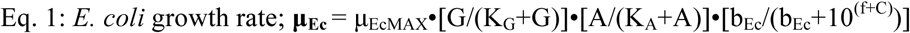

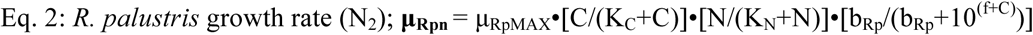

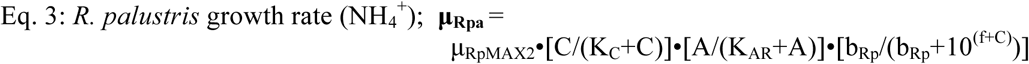

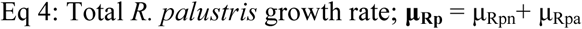

Equations 5-14 were used to describe temporal changes in cell densities and extracellular compounds. Numerical constants in product excretion equations are used to account for molar stoichiometric conversions. Numerical constants used in sigmoidal functions are based on those values that resulted in simulations resembling empirical trends. All R and r parameters are expressed in terms of glucose consumed except for R_A_, which is the amount of NH_4_^+^ produced per *R. palustris* cell (Table S1).

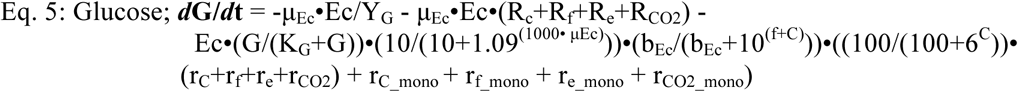

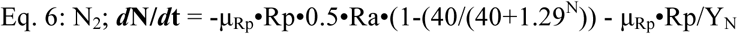

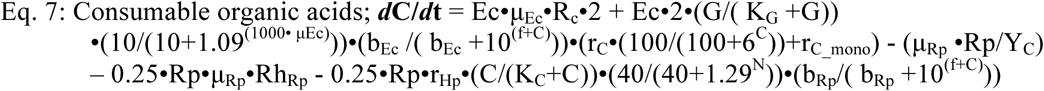

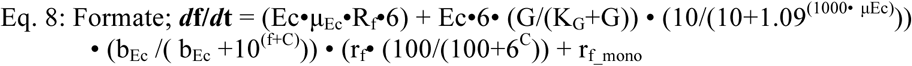

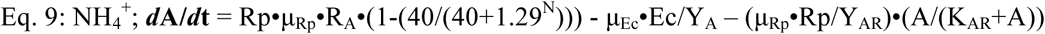

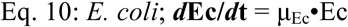

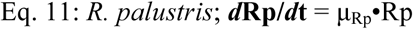

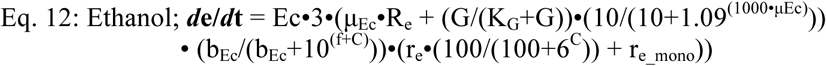

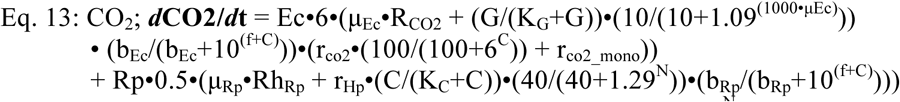

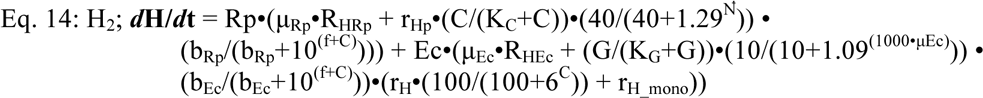

Where,

μ is the specific growth rate of the indicated species (h^-1^).

μ_MAX_ is the maximum specific growth rate of the indicated species (h^-1^).

G, A, C, N, f, e, H and CO2 are the concentrations (mM) of glucose, NH_4_^+^, consumable organic acids, N_2_, formate, ethanol, H_2_, and CO_2_, respectively. All gasses are assumed to be fully dissolved. Consumable organic acids are those that *R. palustris* can consume, namely, lactate (3 carbons), acetate (2 carbons), and succinate (4 carbons). All consumable organic acids were simulated to have three carbons for convenience. Only net accumulation of formate, ethanol, CO_2_ and H_2_ are described in accordance with observed trends.

K is the half saturation constant for the indicated substrate (mM).

Ec and Rp are the cell densities (cells/ml) of *E. coli* and *R. palustris*, respectively.

b is the ability of a species to resist the inhibiting effects of acid (mM).

Y is the *E. coli* or *R. palustris* cell yield from the indicated substrate (cells / μmol glucose). Y values were determined in MDC with the indicated substrate as the limiting nutrient.

R is the fraction of glucose converted into the indicated compound per *E. coli* cell during growth (μmol of glucose / *E. coli* cell), except for R_A_. Values were adjusted to accurately simulate product yields measured in cocultures and in MDC with and without added NH_4_Cl.

R_A_ is the ratio of NH_4_^+^ produced per *R. palustris* cell during growth (μmol / *R. palustris* cell). The default value was based on that which accurately simulated empirical trends.

r is the growth-independent rate of glucose converted into the indicated compound (μmol / cell / h). Default values are based on those which accurately simulated empirical trends in coculture.

r__mono_ is the growth-independent rate of glucose converted into the indicated compound by *E. coli* when consumable organic acids accumulate. Default values are based on linear regression of products accumulated over time in nitrogen-free cell suspensions of *E. coli* (4).

**Table S1.** Default parameter values used in the model unless stated otherwise

| Parameter | Value | Description (Units); Source |
| --- | --- | --- |
| $\mu_{EcMAX}$ | 0.2800 | *E. coli* max growth rate ($h^{-1}$); Monoculture |
| $\mu_{RpMAX}$ | 0.0772 | *R. palustris* max growth rate ($h^{-1}$); Monoculture |
| $\mu_{RpMAX2}$ | 0.0152 | Boost on *R. palustris* growth rate in presence of $NH_4^+$ ($h^{-1}$); Monoculture<sup>a</sup> |
| G | 25 | Glucose (mM) |
| A | 0.00005 | $NH_4^+$ (mM); from initial $(NH_4)_6Mo_7O_{24} \cdot 4H_2O$ concentration |
| C | 0 | Consumable organic acids (those that *R. palustris* was observed to consume: lactate, acetate, and succinate; mM) |
| N | 70 | $N_2$ (assumed to be fully dissolved; mM) |
| f | 0 | Formate (mM) |
| e | 0 | Ethanol (mM) |
| CO2 | 0 | Carbon dioxide (mM) |
| $K_G$ | 0.02 | *E. coli* affinity (Michaelis-Menten constant ($K_m$)) for glucose (mM); (1) |
| $K_C$ | 0.01 | *R. palustris* affinity ($K_m$) for consumable organic acids (mM); Assumed |
| $K_A$ | 0.01 | *E. coli* affinity for $NH_4^+$ (mM); (2) |
| $K_{AR}$ | 0.01 | *R. palustris* affinity for $NH_4^+$ (mM); Assumed<sup>b</sup> |
| $K_N$ | 6 | *R. palustris* affinity ($K_m$) for $N_2$ (mM) |
| Ec | $0.4 \times 10^7$ | *E. coli* cell density (cells / ml) |
| Rp | $3.6 \times 10^7$ | *R. palustris* cell density (cells / ml) |
| $b_{Ec}$ | $10^{43}$ | Resistance of *E. coli* to low pH (mM) |
| $b_{Rp}$ | $10^{32}$ | Resistance of *R. palustris* to low pH (mM) |
| $Y_G$ | $8 \times 10^7$ | Glucose-limited *E. coli* growth yield (cells / μmol glucose); Glucose-limited *E. coli* culture |
| $Y_A$ | $1 \times 10^9$ | $NH_4^+$-limited *E. coli* growth yield (cells / μmol $NH_4^+$); $NH_4^+$-limited *E. coli* culture |
| $Y_C$ | $2.5 \times 10^8$ | Organic acid-limited *R. palustris* growth yield (cells / μmol organic acid); Acetate-limited *R. palustris* culture |
| $Y_N$ | $5 \times 10^8$ | $N_2$-limited *R. palustris* growth yield cells / μmol $N_2$; $N_2$-limited *R. palustris* culture |
| $R_C$ | $1.9 \times 10^{-8}$ | Fraction of glucose converted to organic acids (μmol glucose / cell) |
| $R_f$ | $8 \times 10^{-9}$ | Fraction of glucose converted to formate (μmol glucose / cell) |
| $R_e$ | $4.5 \times 10^{-9}$ | Fraction of glucose converted to ethanol (μmol glucose / cell) |
| $R_{CO2}$ | $5 \times 10^{-10}$ | Fraction of glucose converted to $CO_2$ (μmol glucose / cell) |
| $R_{HRp}$ | $2 \times 10^{-9}$ | *R. palustris* $H_2$ production (μmol $H_2$ / *R. palustris* cell) |
| $R_{HEc}$ | $5 \times 10^{-9}$ | *E. coli* $H_2$ production (μmol $H_2$ / *E. coli* cell) |
| $R_A$ | $0.15 \times 10^{-9}$ | *R. palustris* $NH_4^+$ production (μmol $NH_4^+$ / cell) |
| $r_C$ | $300 \times 10^{-11}$ | *E. coli* specific growth-independent rate of glucose conversion to consumable organic acids (μmol glucose / cell / h) (3) |
| $r_f$ | $47 \times 10^{-11}$ | *E. coli* specific growth-independent rate of glucose conversion to formate (μmol glucose / cell / h) (3) |
| $r_e$ | $15 \times 10^{-11}$ | *E. coli* specific growth-independent rate of glucose conversion to ethanol (μmol glucose / cell / h) (3) |
| $r_{CO2}$ | $2 \times 10^{-11}$ | *E. coli* specific growth-independent rate of glucose conversion to $CO_2$ (μmol glucose / cell / h) (3) |
| $r_H$ | $2 \times 10^{-11}$ | *E. coli* specific growth-independent rate of H<sub>2</sub> production (μmol H<sub>2</sub> / cell / h) (3) |
| $r_{C\_mono}$ | $1.2 \times 10^{-11}$ | *E. coli* specific growth-independent rate of glucose conversion to consumable organic acids when consumable organic acids accumulate (μmol glucose / cell / h); (4) |
| $r_{f\_mono}$ | $0.83 \times 10^{-11}$ | *E. coli* specific growth-independent rate of glucose conversion to formate when consumable organic acids accumulate (μmol glucose / cell / h); (4) |
| $r_{e\_mono}$ | $0.5 \times 10^{-11}$ | *E. coli* specific growth-independent rate of glucose conversion to ethanol when consumable organic acids accumulate (μmol glucose / cell / h); (4) |
| $r_{co2\_mono}$ | $1.3 \times 10^{-11}$ | *E. coli* specific growth-independent rate of glucose conversion to CO<sub>2</sub> when consumable organic acids accumulate (μmol glucose / cell / h); (4) |
| $r_{H\_mono}$ | $0.83 \times 10^{-11}$ | *E. coli* specific growth-independent rate of glucose conversion to H<sub>2</sub> when consumable organic acids accumulate (μmol glucose / cell / h); (4) |
| $r_{Hp}$ | $27 \times 10^{-11}$ | *R. palustris* specific growth-independent rate of H<sub>2</sub> production (μmol H<sub>2</sub> / cell / h) |

<sup>a</sup> Increased growth rate in presence of NH<sub>4</sub><sup>+</sup> versus N<sub>2</sub> based on the difference in experimentally determined growth rates in *R. palustris* monocultures grown with either NH<sub>4</sub><sup>+</sup> or N<sub>2</sub> as a nitrogen source.
<sup>b</sup> K<sub>AR</sub> was assumed to be equivalent to the published *E. coli* K<sub>m</sub> (2) for NH<sub>4</sub><sup>+</sup> (K<sub>A</sub>).

**Table S2.**
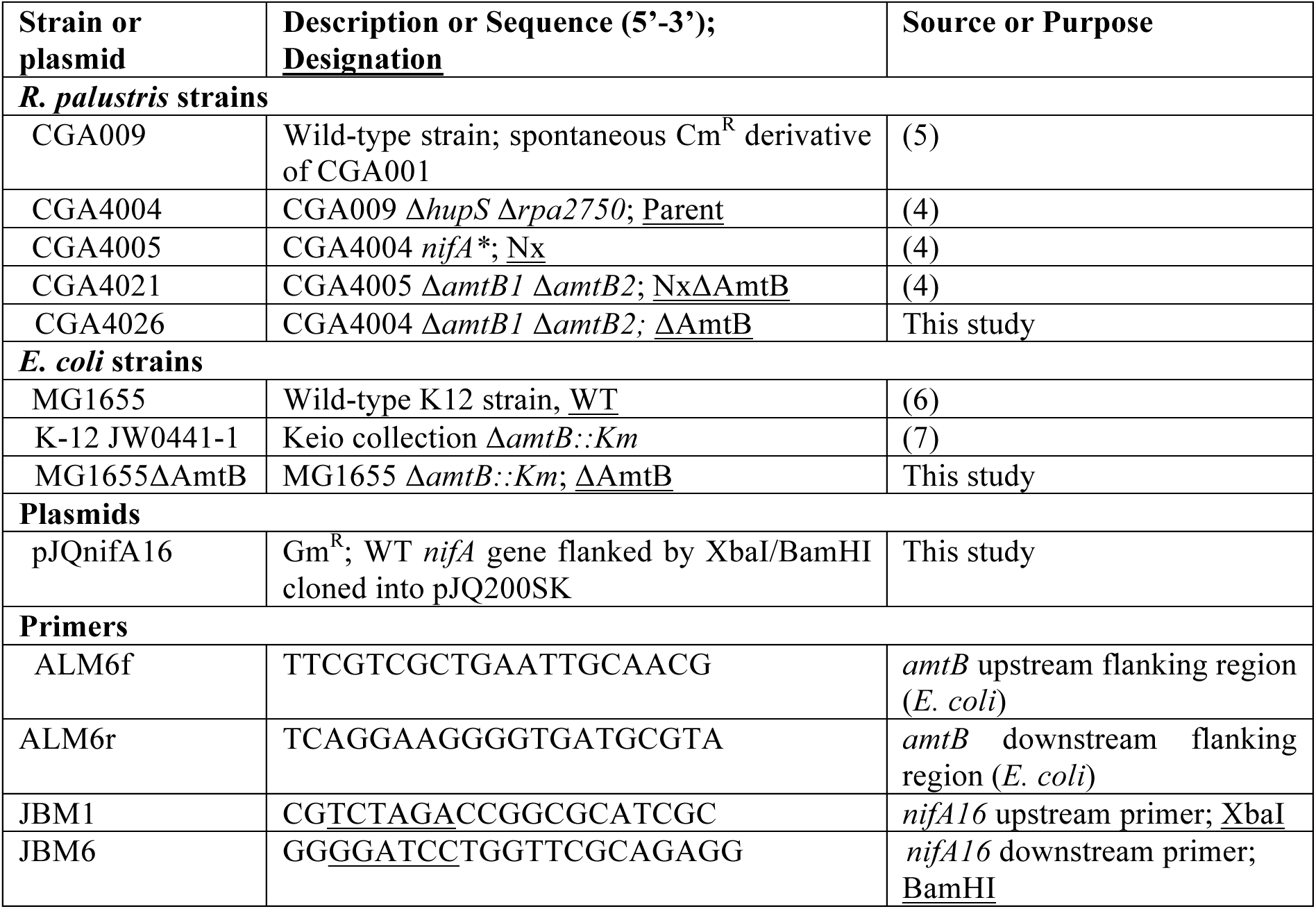
Strains, plasmids, and primers used in this study

### SI Appendix Figures

**Fig. S1.**
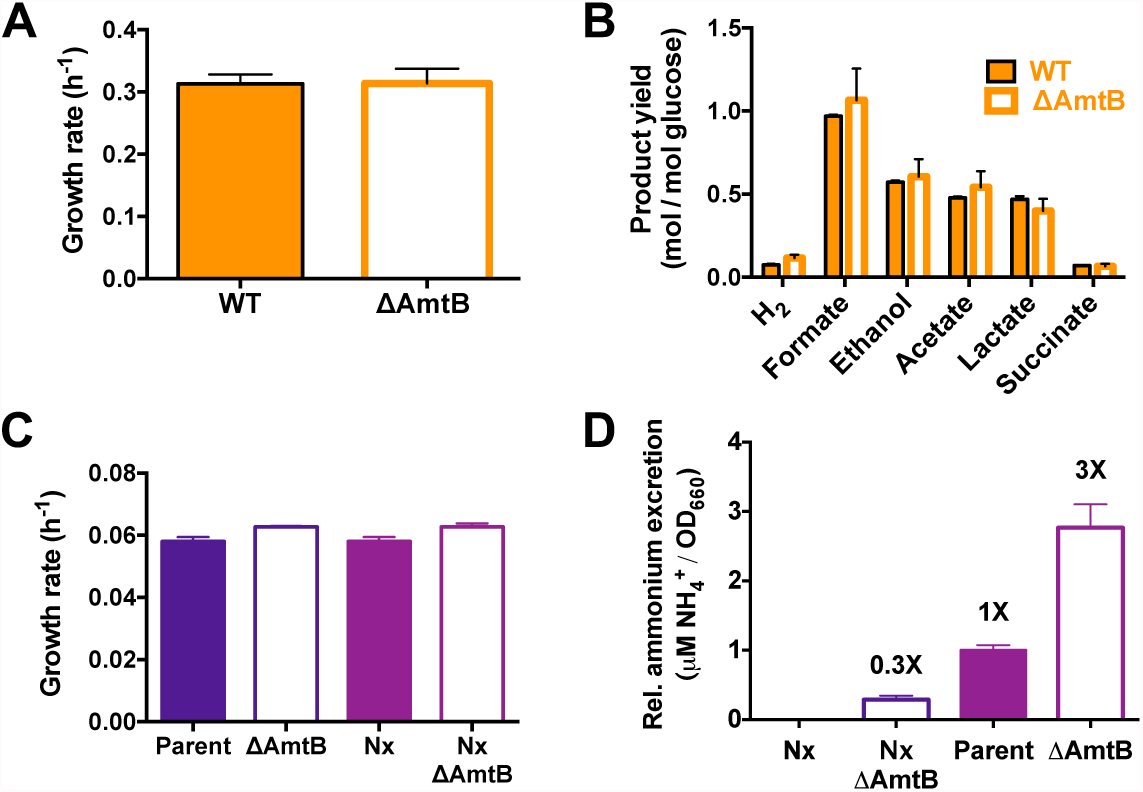
*E. coli* ΔAmtB and *R. palustris* ΔAmtB monoculture growth and metabolic trends. (**A**,**B**) Growth rates (**A**) and fermentation product yields (**B**) from WT *E. coli* (filled) or without (open) *E. coli* ΔAmtB monocultures grown in MDC with 25 mM glucose and 15 mM NH_4_Cl. Fermentation profiles were generated from stationary monocultures. Error bars indicate SD, n=3. (**C**,**D**) Growth curves (**C**) and relative NH_4_^+^ excretion (**D**) of *R. palustris* monocultures grown in MDC with 3 mM sodium acetate and a 100% N_2_ headspace. Error bars indicate SD, n=4.

**Fig. S2.**
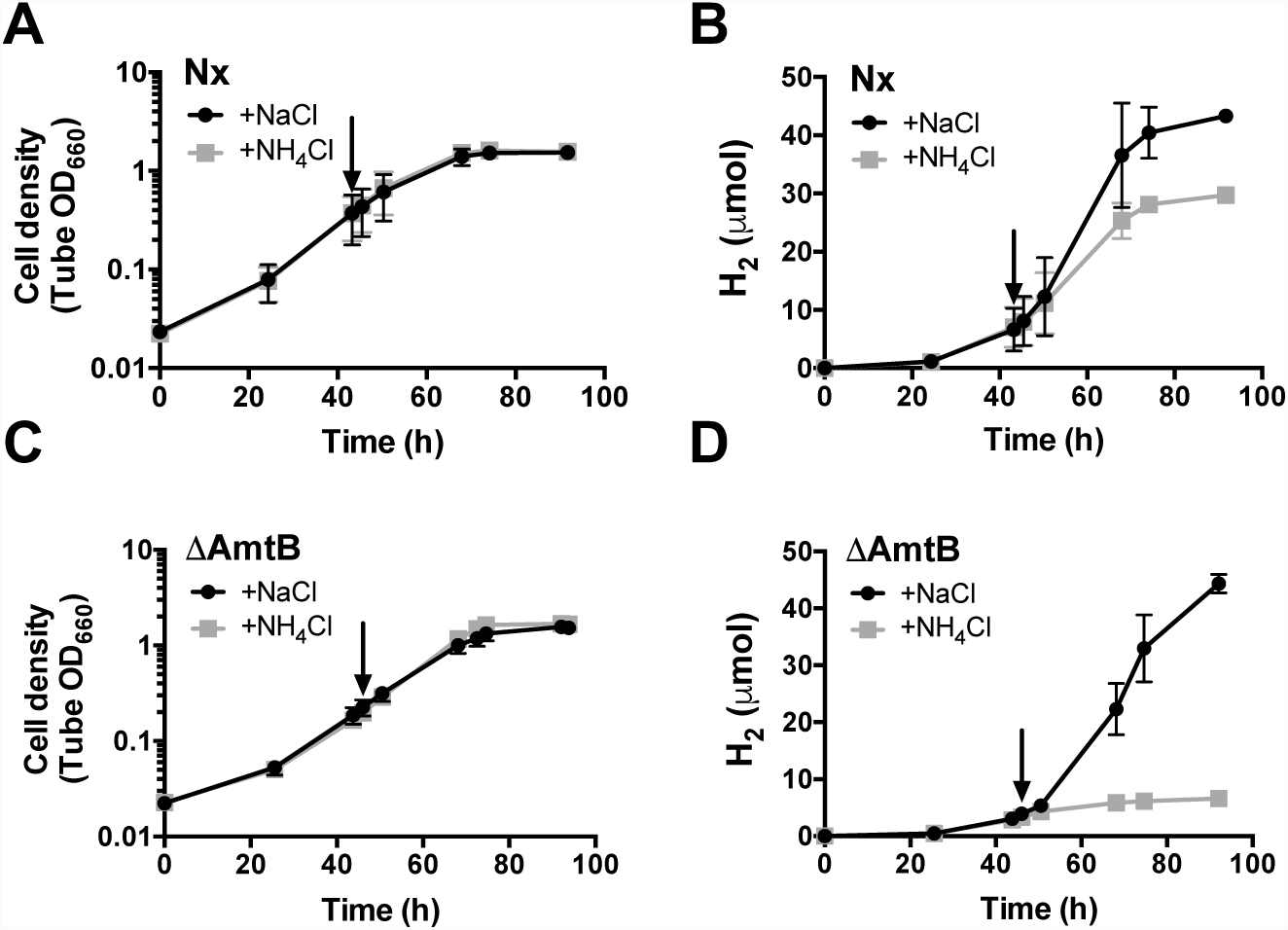
*R. palustris* ΔAmtB responds to NH_4_^+^-induced shutoff of nitrogenase. The effect of either NH4Cl or NaCl on growth **(A**,**C**) and H_2_ production (**B**,**D**) in *R. palustris* Nx or *R. palustris* ΔAmtB monocultures. *R. palustris* monocultures were grown in MDC with 20 mM sodium acetate and a 100% N_2_ headspace until mid-exponential phase and then supplemented with either 15 mM NH_4_Cl or 15 mM NaCl at the time indicated by the arrow.

**Fig. S3.**
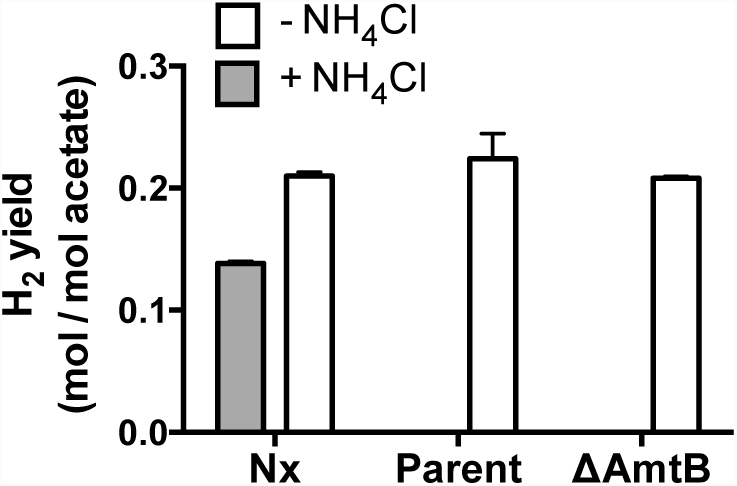
Unlike *R. palustris* Nx, *R. palustris* ΔAmtB does not produce H_2_ when grown with NH_4_^+^. *R. palustris* monocultures were grown in MDC with 20 mM sodium acetate and a 100% N_2_ headspace with (grey) or without (white) 15mM NH_4_Cl. Samples for determining H_2_ yields were taken one week after inoculation, within 24 hours into stationary phase. Error bars indicate SD, n=3.

**Fig. S4.**
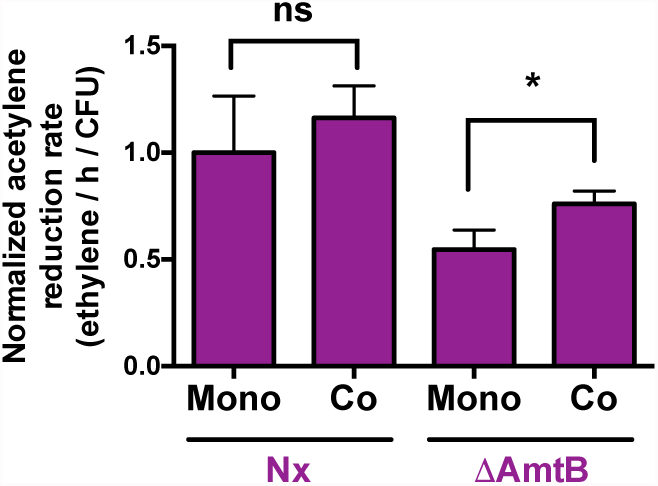
*R. palustris* ΔAmtB nitrogenase activity increases in coculture. Normalized nitrogenase activity of *R. palustris* in monoculture (Mono) or coculture (Co) measured by an acetylene reduction assay. Ethylene levels were divided by total *R. palustris* CFUs in the test tube and then normalized to the *R. palustris* Nx monoculture value. Error bars indicate SD, n=4. *, statistical difference between monoculture and coculture conditions, p < 0.05, determined using multiple two-tailed t-tests; ns, no significant difference.

**Fig. S5.**
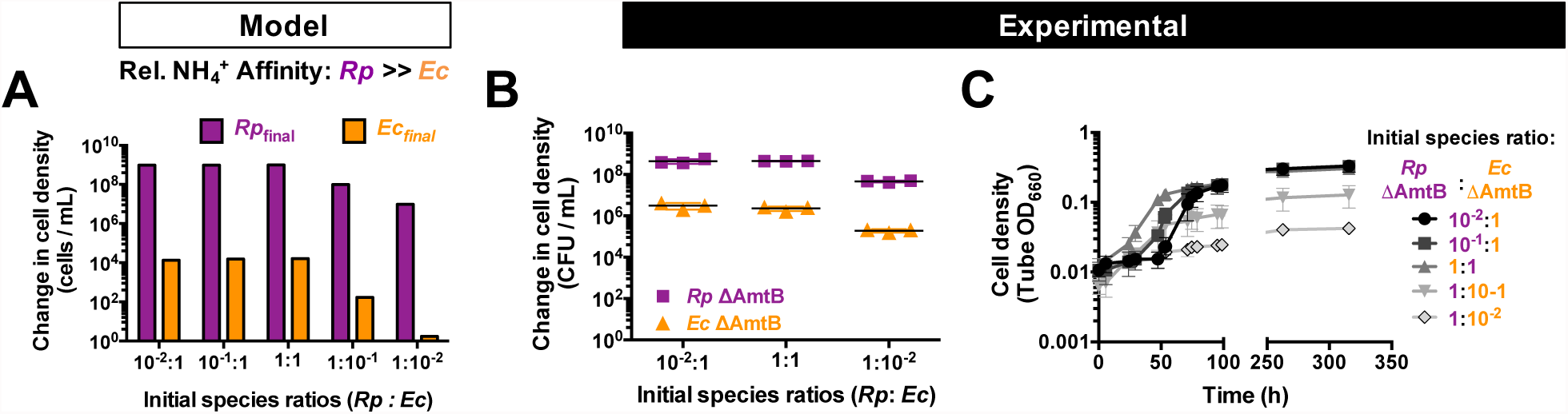
Higher initial cell densities of *E. coli* ΔAmtB can partially compensate for a low *E. coli* NH_4_^+^ affinity. Simulations (**A**) and empirical data (**B**,**C**) showing the effect of initial *E. coli* (*Ec*) cell density on population and coculture growth trends when *E. coli* has a lower affinity for NH_4_^+^ compared to *R. palustris* (*Rp*). (**A**) 300 h batch cultures were simulated with a relative *R. palustris* : *E. coli* (*Rp* : *Ec*) K_m_ value for NH_4_^+^ of 0.001. (**B, C**) Change in cell densities after one week of growth (**B**) and growth curves (**C**) of cocultures inoculated at different species ratios. (**A-C**) A ratio value of 1 represents 2.7 x 10^6^ CFUs/mL, which was experimentally measured from the starting inoculum for both species before diluting to achieve the indicated ratios. Error bars indicate SD, n=3.

